# Improved CAR-T cell activity associated with increased mitochondrial function primed by galactose

**DOI:** 10.1101/2023.09.23.559091

**Authors:** Golda Gross, Suha Alkadieri, Amilia Meir, Orit Itzhaki, Yarden Aharony-Tevet, Shahar Ben Yosef, Angi Zenab, Liat Shbiro, Amos Toren, Tal Yardeni, Elad Jacoby

## Abstract

CD19 CAR-T cells have led to durable remissions in patients with refractory B-cell malignancies; nevertheless, most patients eventually relapse in the long term. Many interventions aimed at improving current products have been reported, with a subset of them focusing on a direct or indirect link to the metabolic state of the CAR-T cells. We assessed clinical products from an ongoing clinical trial utilizing CD19-28z CAR-T cells from patients with acute lymphoblastic leukemia. CAR-T clinical products leading to a complete response had significantly higher mitochondrial function (by oxygen consumption rate) irrespective of mitochondrial content. Next, we replaced the carbon source of the media from glucose to galactose to impact cellular metabolism. Galactose-containing media increased mitochondrial activity in CAR-T cells, and improved *in vitro* efficacy, without any consistent phenotypic change in memory profile. Finally, CAR-T cells produced in galactose-based glucose-free media resulted in increased mitochondrial activity. Using an *in vivo* model of Nalm6 injected mice, galactose-primed CAR-T cells significantly improved leukemia-free survival compared to standard glucose-cultured CAR-T cells. Our results prove the significance of mitochondrial metabolism on CAR-T cell efficacy and suggest a translational pathway to improve clinical products.

## Introduction

Metabolism is a significant determinant of therapeutic success of adoptively transferred T-cell therapy^1^. The metabolic phenotype of T-cells is usually linked to their memory status and exhaustion profile. Naïve and memory T-cells display largely quiescent metabolic profiles relying on mitochondrial ATP as an energy source, while active effector T-cells substantially increase mitochondrial activity and glycolysis^2–4^.

Chimeric antigen receptor T-cell therapy (CAR-T cell) is a highly effective treatment against many forms of B-cell malignancies. Therapeutic failure, either primary (failure to achieve remission) or secondary (relapse), despite antigen expression on the target, remains common, hence the need to improve current products^5–7^. Long term durable remissions are associated with long-term memory CAR-T cells in the product and in patients^7,8^. In a cohort of patients with chronic lymphocytic leukemia, CAR-T cell products with low glycolytic metabolism and low terminal differentiation were associated with improved outcomes^8^, highlighting the link between metabolism and memory and outcome also in CAR-T cell products.

Manipulations affecting CAR-T cell phenotype or function will also affect their metabolism, as CD28 co-stimulation activates glycolysis while 4-1BB enhances mitochondrial respiration^9^. In recent years, many interventions aimed at metabolically altering CAR-T cells have been attempted, in addition to co-stimulation change, mostly involving engineering of key metabolic enzymes (reviewed here^10^). An alternative approach is controlling the nutrient supplementation of CAR-T cells, and by that avoiding additional genetic manipulations. For example, short-chain fatty acids feed into mitochondrial oxidation, and when supplemented can reduce glycolytic reliance, resulting in enhancing the efficacy of adoptively transferred T-cells in therapy models^11^. Other nutrients such as inosine may also have a similar effect^12^.

In this study, glucose in the growth media was replaced by galactose to provide CAR-T cells with a carbon source that is more efficiently utilized through mitochondrial metabolism than by glycolysis. Galactose has been used in many different contexts to force cells to diminish their glycolytic metabolism^13^ and has shown impact on T-cell therapy^14^. The current study shows that CAR-T cell products with improved mitochondrial function lead to significantly better antitumor activity *in-vitro* and *in-vivo*, and provides a straightforward and cost-effective approach to improve mitochondrial function in CAR-T cells using galactose supplementation.

## Methods

### Ethics and clinical samples

Patient’s CAR-T cell products were obtained as part of a clinical trial approved by the Sheba Medical Center IRB and the Israeli Ministry of Health (NCT02772198). Production details of clinical CAR-T cells were previously reported^15^. All animal experiments were approved by Institutional Ethical Review Process Committees and were performed under Israel Institutional Animal care and use committee approval (1308/21).

### CAR-T cell production

CAR-T cell production was performed as previously described^16^. Briefly, peripheral-blood mononuclear cells were activated using 50ng/mL OKT3, and transduced with an MSGV CD19-CAR plasmid, containing a CD19-targeting CAR (FMC63-CD28-CD3ζ). CAR-T cells were then grown for 7-11 days.

### Cell culture and cell lines

CAR-T cells were cultured in glucose or galactose-based media (10mM for both) containing 100IU/ml IL-2 and sodium pyruvate (1mM for the latter. Co-culture experiments were performed without additional IL-2. CD19 positive target cell lines Nalm6 (Acute lymphoblastic leukemia, ALL), REH (ALL) and Toledo (Lymphoma) were grown in standard culture conditions^16^.

### Flow cytometry

All flow-cytometry readings were performed on a Gallios flow cytometer (Beckman Coulter). The antibodies used are reported in Supplementary Table 1. Data was analyzed using the FlowJo software.

### Mitochondrial assays

#### Mitochondrial mass and membrane potential

were assessed by MitoTracker Green (MTG, Invitrogen) and tetramethylrhodamine methyl ester perchlorate (TMRE, Abcam), respectively. CAR-T cells (300,000) were stained according to manufacturer instructions and subjected to flow-cytometry.

#### Mitochondrial DNA (mtDNA) copy number

was quantified on isolated DNA (Qiagen). qPCR was performed for mitochondrial tRNA^leu^ in FastSYBR Green Master Mix and the nuclear gene β2-microglobulin (β2M) on a StepOnePlus Real Time PCR System. Primers are reported in Supplementary Table 2. mtDNA copy number was quantified by calculating delta CT of tRNA^leu^ to β2M.

#### ATP quantification

was done using the CellTiter Glo Kit (Promega).

#### Mitochondrial oxygen consumption rate (OCR)

was measured in CAR-T cells using the Seahorse XFe96 Instrument (Seahorse Bioscience). 200,000 cells per well were seeded on poly-D-lysine-coated plates on the same day of the experiment according to manufacturer instructions. The following mitochondrial respiration parameters were measured: (i) basal respiration; (ii) leak respiration (Oligomycin A [Oligo]); (iii-iv) maximal respiration (carbonyl cyanide 3-chlorophenylhydrazone [CCCP, C2759, Sigma] added twice to fully reach mitochondrial reserve capacity); and (v) non-mitochondrial O_2_ consumption (rotenone +antimycin A). For analysis, each well was normalized for the protein content, using Pierce™ BCA Protein Assay Kits

#### CAR-T cell activation assays

CAR-T-cells were plated in their respective media and activated once using irradiated target cells at a 1:2 effector to target (E:T) ratio or with live Nalm6 at a 2:1 E:T ratio, and left to grow for 3 days. In the long activation assays, this process was repeated 3-4 times over a total period of 2 weeks.

#### CAR-T cell long cytotoxicity assay

CAR-T cells were plated with targets at three different E:T ratios, with a control condition of target cells only. Co-culture wells were monitored for a week and media was changed every 2-3 days. After a co-culture period of 7 days, the cells were counted and analyzed by flow cytometry to check for cell survival and identity.

#### In vivo assay

NOD^-^SCID^-^IL-2Rγ^-^ (NSG) mice were purchased from the Jackson laboratories. Six to ten-week-old female NSG mice were tail-vein injected with 0.5x10^6^ Nalm6 cells, and after 3 days treated IV with 2x10^6^ CAR-T cells or untransduced T-cells (controls). Peripheral blood was collected 3 weeks later and analyzed for Nalm6, human CD45+ and CAR+ cells. Mice were monitored at least twice weekly for signs of leukemia, graft-vs-host disease, and survival. Moribund mice were sacrificed per institutional guidelines.

#### Statistical testing

All statistical analysis was done on GraphPad Prism 9 software. Standard deviations were calculated from technical repeats, and standard error of mean was plotted with biologic repeats. Where applicable, paired t-test or ANOVA were used to obtain p-values of significance with pairing within each donor across conditions. Survival was analyzed by the Kaplan-Meier method.

## Results

### Clinical product mitochondrial function analysis

We analyzed CAR-T cell products from patients with ALL enrolled on clinical trial NCT02772198^15,17^ for mitochondrial function and content. CAR-T cell products from patients that failed to respond to therapy had significantly decreased basal and maximal oxygen consumption rates (Fig. 1A-C). Testing on additional products confirmed these observations (supplementary Fig. 1). In addition, we measured the mitochondrial activity of these CAR-T cells by TMRE (Fig. 1D) and mitochondrial mass using both MTG staining by flow-cytometry (Fig. 1E) and qPCR for mtDNA copy numbers (Fig. 1F). The per-sample ratio of TMRE/MTG mean fluorescent index (MFI) was taken as an indicator of activity per mitochondrion. Mitochondrial mass in the non-responders’ products was not decreased (Fig. 1E-F) compared to products of responders. Calculation of the per-sample ratio of TMRE/MTG MFI, representing activity per mitochondrion, demonstrated a trend of increased ratio in responders compared to non-responders (p=0.07, Fig. 1G). Previously, increased mitochondrial activity was associated with increased populations of memory or stem-cell memory CAR T-cells^9,18^. We quantified by flow cytometry memory markers CD45RA and CD62L, and found that products leading to a response tended to show increased effector-memory and effector populations, although this did not reach statistical significance (supplementary Fig. 1). Our data suggests that CAR-T cell products with increased mitochondrial function per mitochondrion led to better outcomes.

**Fig. 1:**
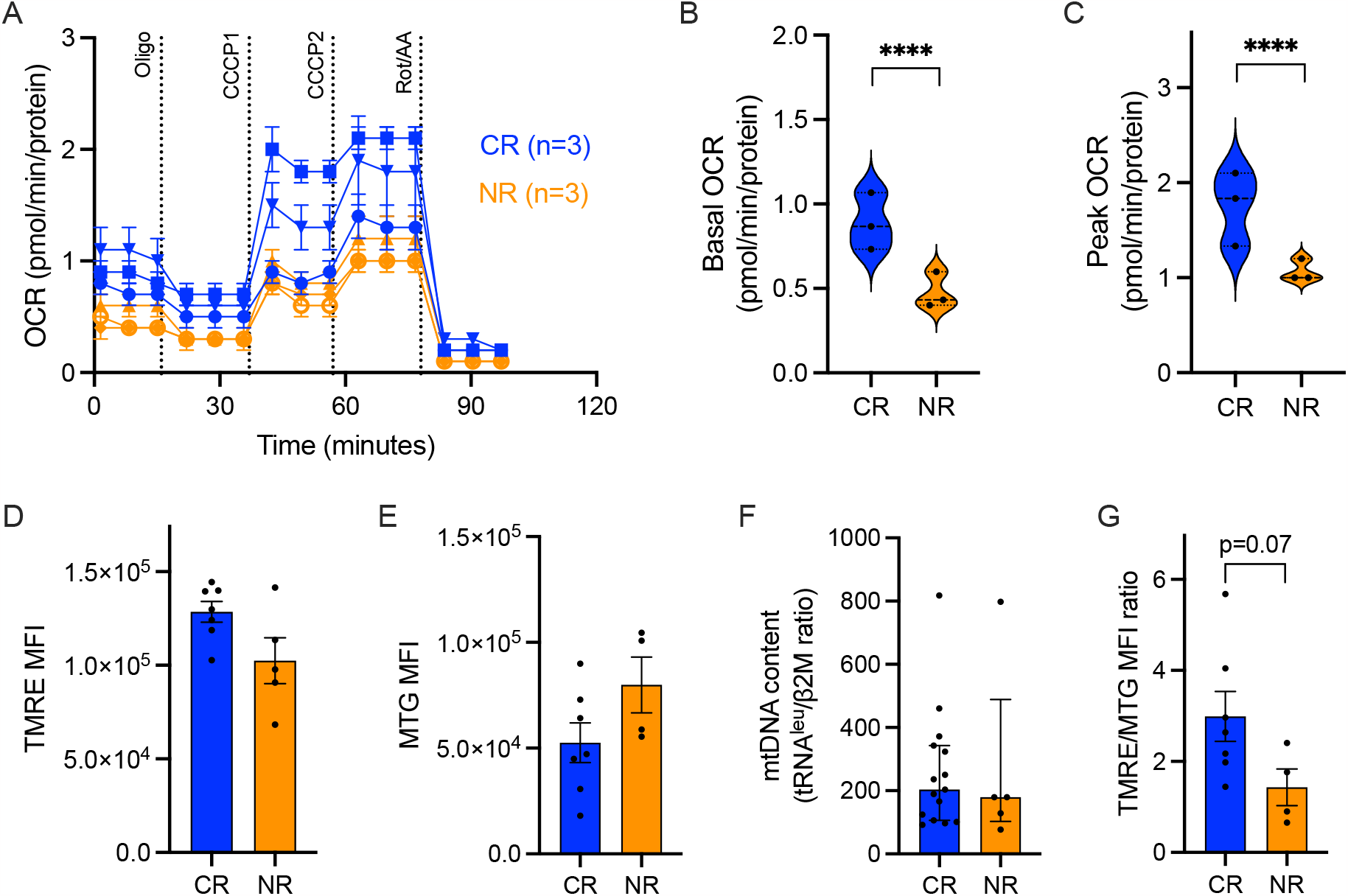
Mitochondrial parameters of clinical CAR-T cell products of six pediatric ALL patients. (A) Oxygen Consumption Rate (OCR) measurements were obtained over time (minutes) using the Seahorse XFe96 analyzer and normalized to the amount of protein per well. Using the mitochondrial stress assay, we tested basal mitochondrial respiration, mitochondrial proton leak by adding the ATP synthase inhibitor Oligomycin A (1.5μM), followed by the mitochondrial uncoupler cyanide 3-chlorophenylhydrazone (CCCP) twice, (4μM total) for measuring maximal respiration, and Rotenone (0.5μM) together with Antimycin A (0.5μM) for non-mitochondrial respiration. (B) Basal mitochondrial OCR and (C) maximal respiration. (D) Mitochondrial activity measured by mean fluorescent index (MFI) of TMRE staining by flow cytometry. (E) Mitochondrial mass measured by MFI of MitoTracker Green (MTG) staining by flow cytometry; (F) Mitochondrial content measured by mtDNA copy number, by qPCR of tRNA^leu^ and β2-microglobulin (β2M), and (G) Ratio of the MFI of TMRE to the MFI of MTG per sample. Complete response patients, CR (blue, N=3), non-responding patients, NR (orange, n=3). P value ****<0.0001.

### Galactose enhances mitochondrial activity in CAR-T cells

Galactose has been shown to impact on T-cell function^2,14^ and to affect T-cell metabolism by decreasing glycolysis and increasing dependence on mitochondrial activity^11^. Here, we verified the metabolic changes following culture in galactose (vs. glucose) in CD19 CAR-T cells. The addition of Oligo, an inhibitor of the mitochondrial ATP synthase, proved the dependence of galactose-cultured CAR-T cells on mitochondrial metabolism, by causing a significant drop in ATP levels (Fig. 2A) in both resting and activated CAR-T cells, and an increase in cell death (Fig. 2B), only when cultured in galactose.

**Fig. 2:**
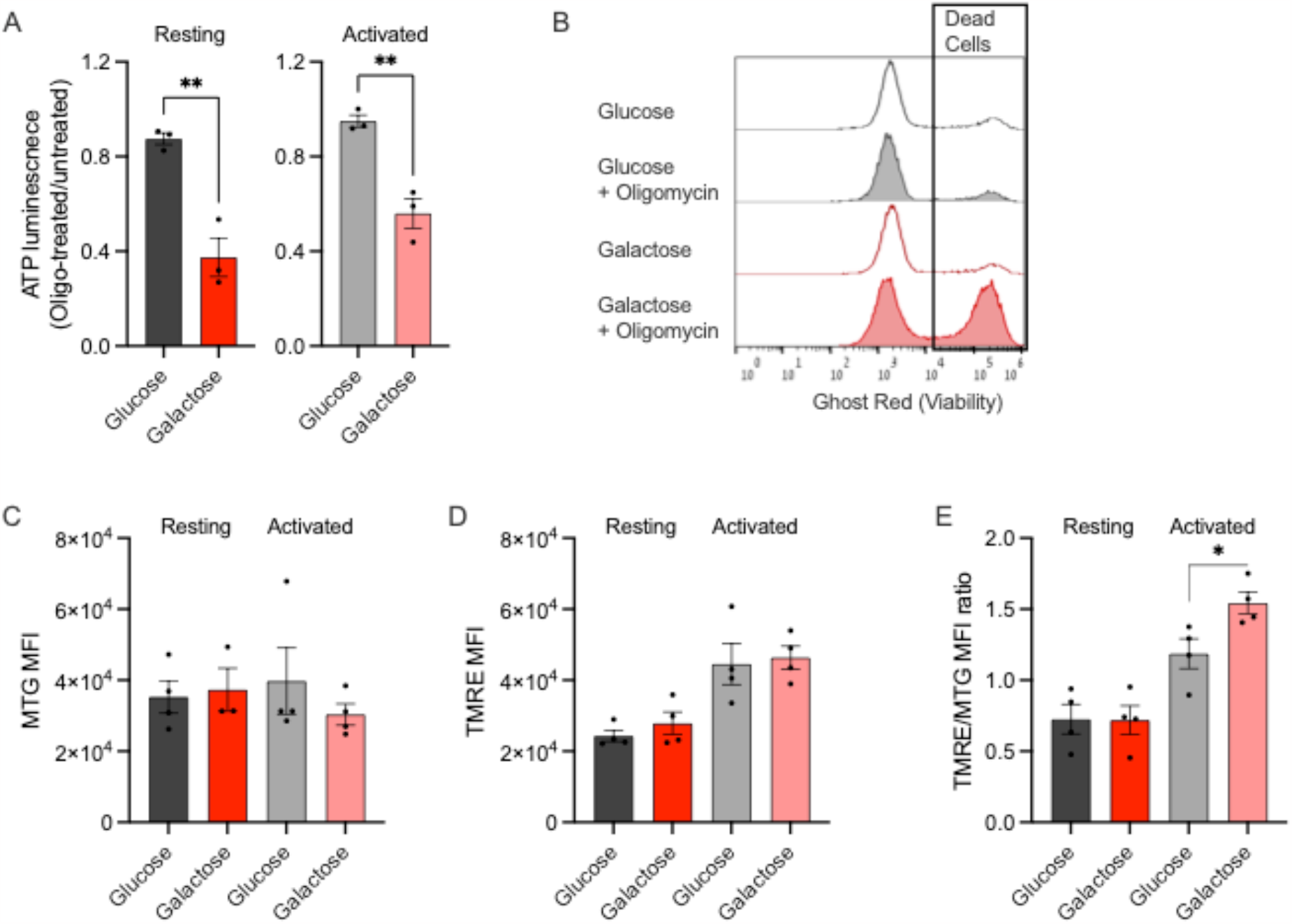
CD19 CAR-T cell metabolism and activity in glucose and galactose. (A) ATP content was assessed by luminescence assay in Oligomycin A-treated (1uM) resting (left) or activated (right) CAR-T cells vs untreated cells in glucose (grey) or galactose (red). (B) Ghost red staining indicating cell death in glucose and galactose cultured CAR-T cells with or without Oligomycin A, read by flow cytometry. (C-E) Resting and Nalm6-activated CAR-T cells were stained with TMRE or MTG followed by flow cytometry. Mean fluorescence index (MFI) of MTG (D), TMRE (E) and the per-sample ratio of TMRE/MTG MFI are shown in all conditions. P value *<0.05, **<0.01.

CAR-T cell activation results in increased mitochondrial activity, as measured by TMRE MFI, regardless of mitochondrial content measured by MTG MFI (supplementary Fig. 2). We evaluated the mitochondrial content by flow cytometry for MTG, and it did not change between CAR-T cells in either media or following activation (Fig. 2C). In activated cells in both culture media the mitochondrial membrane potential increased as expected (Fig. 2D). However, activated galactose-cultured cells displayed a higher TMRE/MTG MFI ratio than when activated in glucose, suggesting higher activity per mitochondrion. Altogether, as expected, we show a shift towards mitochondrial metabolism in galactose-cultured CAR-T cells.

### Improved CAR-T cell activity due to galactose supplementation

To test whether galactose-mediated metabolic changes affect CAR-T cell function, we co-cultured CAR-T cells with the CD19-positive Nalm6 or REH cell lines across a range of effector to target (E:T) ratios for a week in either media. Growth of target cells was monitored in glucose and galactose, and was similar for REH cells in both media, but slower for Nalm6 in galactose than in glucose (Supplementary Fig. 3). At an E:T ratio of 1:8, CAR-T cells efficiently killed both targets regardless of the culture conditions. However, at E:T ratios of 1:16 and 1:32, there was a significant killing advantage for galactose co-cultured cells (Fig. 3A-B), against both Nalm6 and REH targets.

**Fig. 3:**
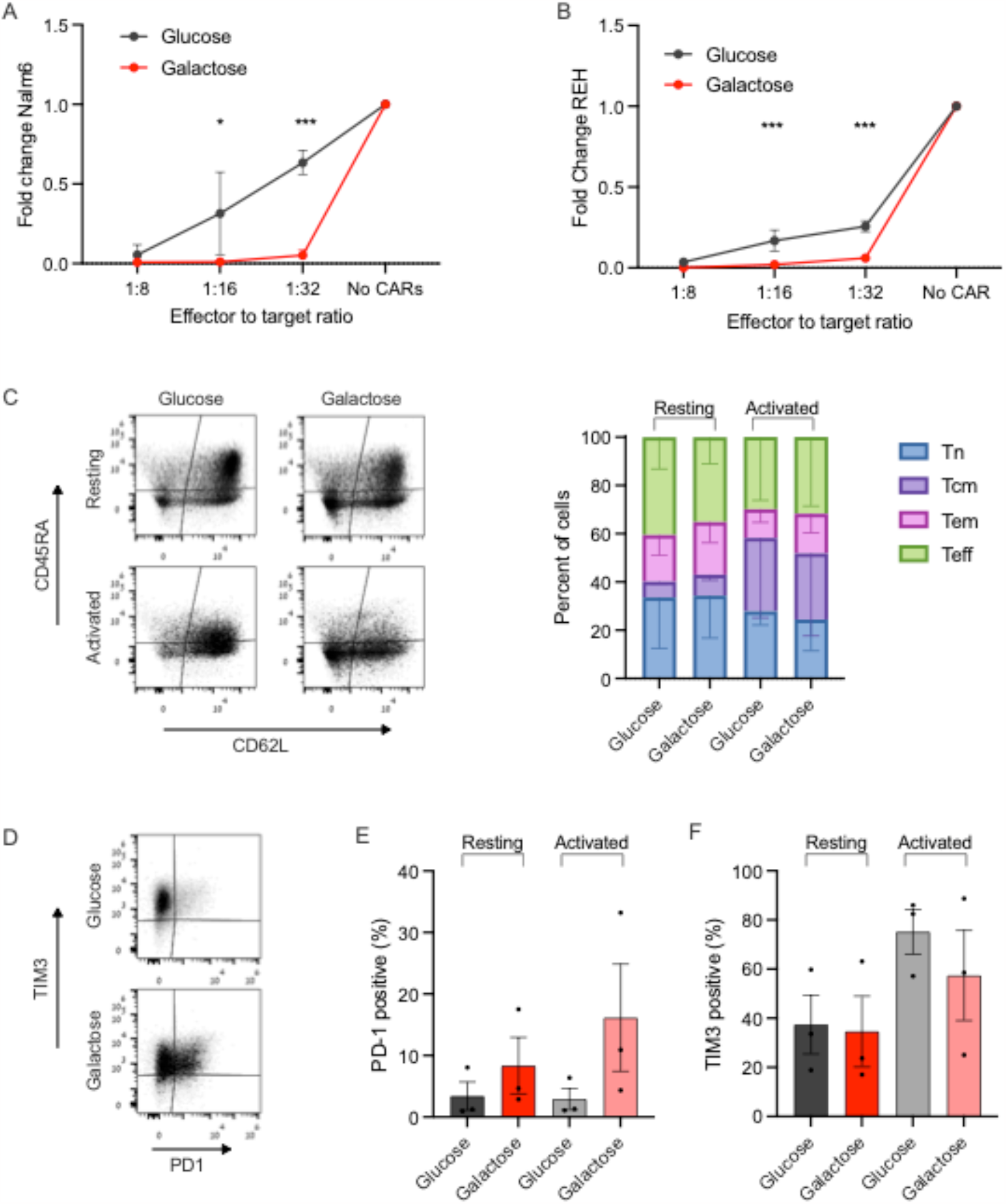
Functional properties of CAR-T cells are impacted by galactose. (A-B) Nalm6 (A) and REH (B) leukemia cells were co-cultured with CAR-T cells for a week under effector to target ratios of 1:8, 1:16 and 1:32, in either glucose or galactose. Fold change of Nalm6 cells (A) or REH cells (B) compared to their growth in the same media without CAR-T cells is shown. Target cells were identified by flow cytometry as CD10+. (C-F) Glucose or galactose-cultured CAR-T cells, either resting or activated with CD19+ targets, were subjected to flow cytometry analysis for memory subsets (C), PD-1 (D-E) and TIM3 (D,F) expression. Memory subsets were based on CD45RA and CD62L expression, and reported as naïve (Tn, blue), central memory (Tcm, violet), effector memory (Tem, pink) and effector (Teff, green). P value *< 0.05 ***<0.001.

Mitochondrial function is associated with memory T-cell subsets^3,9^. However, here CAR-T cells grown and repeatedly activated in glucose and galactose for 14 days had similar proportions of naïve, central memory, effector-memory or effector cells (Fig. 3C), suggesting that the metabolic and functional changes we observed are independent of the memory status of the cells. PD-1 expression was higher in galactose cultured cells, though this did not reach statistical significance. TIM3 expression did not change significantly between CAR-T cells cultured in glucose vs. galactose, in resting or in activated CAR-T cells (Fig. 3D-F). Thus, improved CAR-T cell functionality was seen in galactose, was related to mitochondrial function and independent of cellular exhaustion markers or memory states.

### Priming CAR-T cells in galactose is functionally beneficial

Our data demonstrated improved efficacy and mitochondrial activity of CAR-T cells cultured in galactose. To translate these findings into the clinical arena we opted to assess whether this modification can be introduced throughout the production process. Therefore, we activated and transduced T-cells from healthy donors in either glucose or galactose. The median fold change in cell numbers during the production of CAR-T cells was significantly higher in glucose-grown CAR-T cells (GluCARs) compared to galactose-grown CAR-T cells (GalaCARs, Fig. 4A and supplementary Fig. 4). Along with the slower growth of GalaCARs, their transduction efficacy was significantly lower than that of the GluCARs (Fig. 4B). The final GalaCAR product had a higher proportion of PD-1 positive cells, similar TIM3 expression and a similar memory T-cell subsets compared to the standard GluCAR product (Fig. 4C-E). GalaCARs also had higher basal and peak OCR compared to GluCARs (Fig. 4F-H). Flow-cytometry analysis for TMRE and mtDNA content did not present any differences between products at rest (Fig. 4I-J). However, once activated, GalaCARs had a greater capacity of increasing mitochondrial activity as measured by TMRE/MTG MFI ratio (supplementary Fig. 5).

**Fig. 4:**
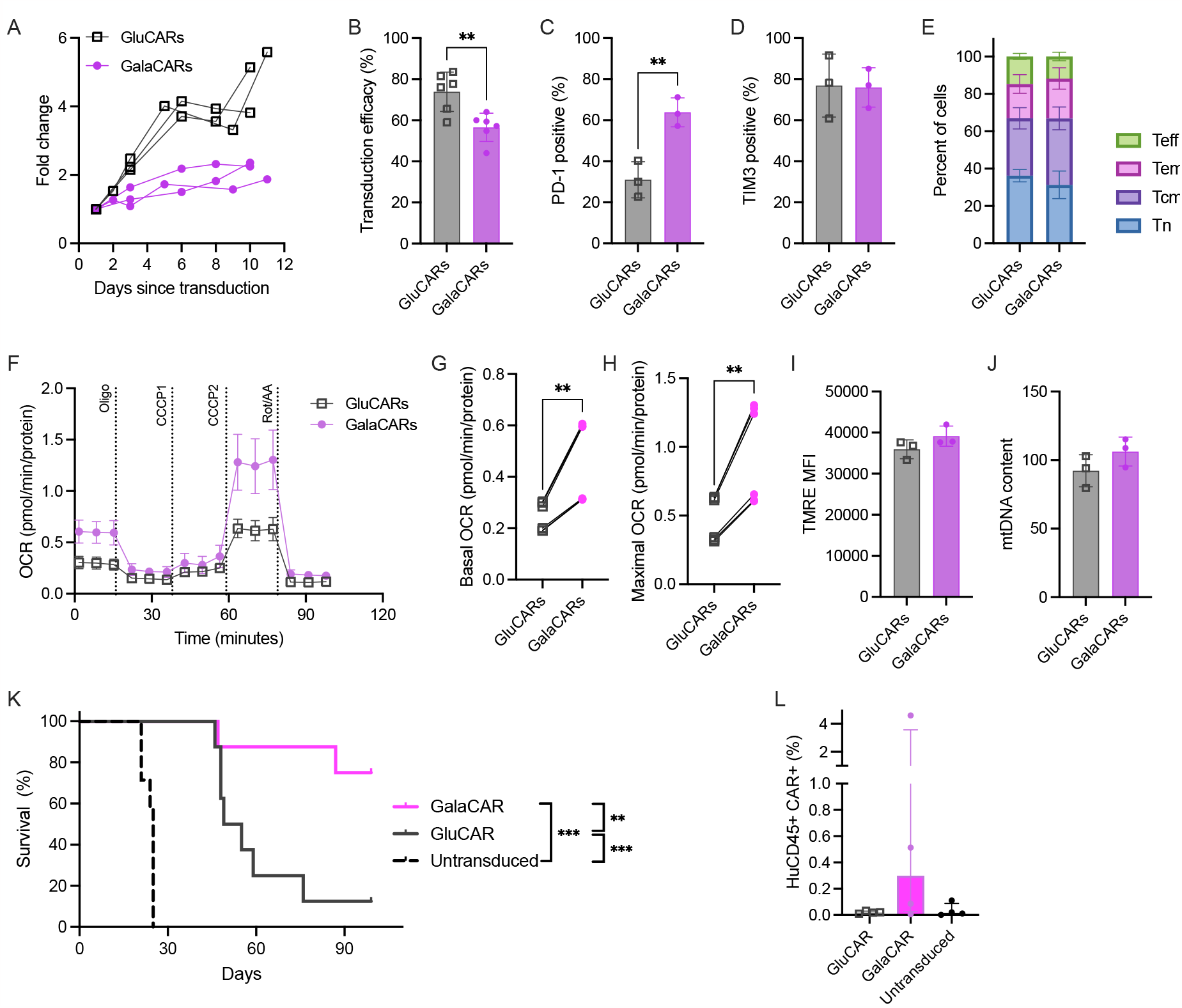
CD19 CAR-T cells were produced in glucose (GluCARs, grey) or galactose (GalaCARs, magenta). (A) Growth of the T-cells in culture during production starting at transduction. (B-E) GluCARs and GalaCARs were assessed by flow cytometry for (B) transduction efficacy; (C) PD-1 expression; (D) TIM3 expression; and (E) memory profile [naïve (Tn, blue), central memory (Tcm, violet), effector memory (Tem, pink) and effector (Teff, green)]. (F-J) Mitochondrial parameters of GluCARs and GalaCARs were assessed by Seahorse XF96. Oxygen consumption rate (OCR) measured in cells from a single donor is shown in (F). Basal OCR (G) and maximal respiration (H) on multiple products from 2 donors are shown. (I) Mean fluorescent index of TMRE is shown in GluCARs (grey) and GalaCARs (magenta). (J) mtDNA content by RT-qPCR for mitochondrial gene tRNA^leu^ and nuclear gene β2-microglobulin. (K) Survival curves for NSG mice injected with Nalm6 cells that were treated with untransduced T-cells, GluCARs or GalaCARs (n=8 per group, 2 independent experiments). (L) Peripheral blood was drawn from treated mice on day 14 after CAR-T cell or mock T-cell treatment and analyzed for human CD45 and CAR expression. Percent of cells in blood is shown. Experiments done on 3-6 independent donors. P value **<0.01. ***<0.001

Demonstration of improved mitochondrial capacity in GalaCARs led us to functionally assess them vs. the standard-grown GluCARs under physiological conditions. Thus, an *in vivo* efficacy assay was carried out, in which Nalm6-bearing NSG mice were treated with either GluCARs or GalaCARs, with untransduced T-cells serving as controls. Both CAR-T products provided long-term benefit over untransduced T-cells, but the survival of GalaCAR treated mice was significantly improved compared to the standard GluCAR treated mice (Fig. 4K). This was associated with improved durability of the CAR-T cells *in vivo* (Fig. 4L). These data confirm that priming the T-cells by replacing glucose with galactose increases mitochondrial activity and results in a superior CAR-T product.

## Discussion

In recent years, numerous studies have revealed that cellular energy production is tightly connected to cell function and phenotype. T-cells have different metabolic strategies depending on their life cycle, and this has influenced the field of adoptive cell therapy in the direction of metabolic manipulation of CD8 T-cells to improve their antitumor function^10^. Successful attempts included glucose restriction^19^, glycolytic inhibition^20^, inhibition of the mitochondrial pyruvate carrier^21^ and inosine supplementation^12^ among others, which have all led to better antitumor responses. These studies suggest that the benefits of pre-conditioning T-cells during their expansion phase are maintained even after the adoptive cell product has been transferred to the patient. Moreover, they validate that improving mitochondrial fitness is key to ensuring CD8 T-cell efficacy *in vivo*. Still, differences in phenotype, signaling and activation pathways of CAR-T cells compared to unmanipulated T-cells^22^ would require confirming such findings in the CAR field.

Our initial results showed that CAR-T cell products with higher mitochondrial respiration, spare respiratory capacity and membrane potential were associated with response to treatment, without showing an increase in mitochondrial mass. These results prompted us to attempt metabolic conditioning that would enhance CAR-T cell mitochondrial activity.

Since CAR transduction is already a major genetic alteration to the patient cells, we opted for an alternative method of enhancing mitochondrial activity consisting of growing the cells in media in which galactose replaced glucose as a carbon source. Growing cells in galactose is now considered a bona fide method to force cells to rely on mitochondria as an energy source^2,23,24^. In CAR-T cells, this shift to mitochondrial metabolism occurred without changes in memory marker expression, and led to increased efficacy *in vitro* even though PD-1 was increased. Previous studies demonstrated the association of improved mitochondrial function in CAR-T cell products of patients with chronic lymphocytic leukemia (CLL) with response^8,25^ and in preclinical models aimed at improving T-cell functionality^6,15^. However, products leading to response in these studies had an increased proportion of memory cells and reduced exhaustion profiles, in addition to high mitochondrial function^8,25^. Here we verified that mitochondrial activity, both in patient products and in our *in vitro* work, is an independent factor for CAR-T cell quality. The CAR structure, containing CD28 vs 4-1BB costimulation, and the source of the cells – from patients with ALL and not CLL, may help explain the different findings.

Additionally, we confirmed that the production of CAR-T cells in galactose-based media led to more efficient products *in vivo*. A similar approach of altering the media during CAR-T cell production was recently published using inosine as a carbon source. Inosine-grown CD8+ T-cells had superior activity; however this was demonstrated in tumor models that cannot compete for this substrate^12^. Our results showed benefit of galactose-cultured CAR-T cells irrespective of the targets’ growth under the same conditions, and *in vivo* in a physiological environment. We suggest that the enhancement of mitochondrial metabolism was the source of the improved effect of GalaCARs reported here. An alternative explanation may be the mild inhibition of growth in culture. Recently, use of dasatinib or other signaling inhibitors during CAR-T cell production led to signaling inhibition and increased efficacy^26^.

The study’s main limitation is the use of only CD28-costimulated CAR-T cells, both in our clinical products and preclinical experiments. CD28-costimulation is associated with a higher glycolysis and lower mitochondrial function compared to 4-1BB costimulation^9^. Still, approved products for lymphoma and ALL contain CD28-costimulation, and it is therefore critical to improve current production platforms.

In summary, we show that increased activity per mitochondrion is a feature of better CAR-T cell products. We show that culturing CAR-T cells in galactose shifts their metabolism towards mitochondrial activation, leading to superior *in vitro* and *in vivo* activity. Translation of these findings may lead to increased response rates of CAR-T cell treated patients.

## Acknowledgements

The authors thank the production team at the Ella Institute of Immunooncology who produced the clinical CAR-T cells. This work was carried out in partial requirements for GG and SA’s MSc degree (Faculty of Medicine, Tel Aviv University).

This work was funded by the Dotan center for hematologic malignancies grant (EJ) and NIH grant 5R01CA259635 (TY).

## Authors’ contributions

GG, TY and EJ conceived the initial study plan. GG, SA, AM, YAT, SBY AZ, LS, TY and EJ performed experiments and analyzed the data. AT, TY and EJ supervised the work. GG, TY and EJ wrote the manuscript draft. All authors approved the manuscript.

## Competing Interests statement

The authors report no financial conflict of interests.

## Data availability

All data is available upon request

## Notes

<u>Competing Interests statement</u>: The authors report no financial conflict of interests. This work was funded by the Dotan center for hematologic malignancies grant (EJ) and NIH grant 5R01CA259635 (TY).

### Competing Interest Statement

The authors have declared no competing interest.

## References

1. Kishton RJ, Sukumar M, Restifo NP. Metabolic Regulation of T Cell Longevity and Function in Tumor Immunotherapy. Cell Metab. 2017;26(1):94–109. doi:10.1016/j.cmet.2017.06.016

2. Chang CH, Curtis JD, Maggi LB, et al. Posttranscriptional control of T cell effector function by aerobic glycolysis. Cell. 2013;153(6):1239. doi:10.1016/j.cell.2013.05.016

3. Kondo T, Ando M, Nagai N, et al. The NOTCH–FOXM1 axis plays a key role in mitochondrial biogenesis in the induction of human stem cell memory–like CAR-T cells. Cancer Res. 2020;80(3):471–483. doi:10.1158/0008-5472.CAN-19-1196

4. Vardhana SA, Hwee MA, Berisa M, et al. Impaired mitochondrial oxidative phosphorylation limits the self-renewal of T cells exposed to persistent antigen. Nat Immunol. 2020;21(9):1022–1033. doi:10.1038/s41590-020-0725-2

5. Labanieh L, Mackall CL. CAR immune cells: design principles, resistance and the next generation. Nature. 2023;614(7949):635–648. doi:10.1038/s41586-023-05707-3

6. Shah NN, Fry TJ. Mechanisms of resistance to CAR T cell therapy. Nat Rev Clin Oncol. 2019;16(6):372–385. doi:10.1038/s41571-019-0184-6

7. Cappell KM, Kochenderfer JN. Long-term outcomes following CAR T cell therapy: what we know so far. Nat Rev Clin Oncol 2023. April 2023:1–13. doi:10.1038/s41571-023-00754-1

8. Fraietta JA, Lacey SF, Orlando EJ, et al. Determinants of response and resistance to CD19 chimeric antigen receptor (CAR) T cell therapy of chronic lymphocytic leukemia. Nat Med. 2018;24(5):563–571. doi:10.1038/s41591-018-0010-1

9. Kawalekar OU, O’Connor RS, Fraietta JA, et al. Distinct Signaling of Coreceptors Regulates Specific Metabolism Pathways and Impacts Memory Development in CAR T Cells. Immunity. 2016;44(2):380–390. doi:10.1016/j.immuni.2016.01.021

10. Peng J-J, Wang L, Li Z, Ku C-L, Ho P-C. Metabolic challenges and interventions in CAR T cell therapy. Sci Immunol. 2023;8(82):eabq3016. doi:10.1126/sciimmunol.abq3016

11. Luu M, Riester Z, Baldrich A, et al. Microbial short-chain fatty acids modulate CD8+ T cell responses and improve adoptive immunotherapy for cancer. Nat Commun. 2021;12(1):1–12. doi:10.1038/s41467-021-24331-1

12. Wang T, Gnanaprakasam JNR, Chen X, et al. Inosine is an alternative carbon source for CD8+-T-cell function under glucose restriction. Nat Metab. 2020;2(7):635–647. doi:10.1038/s42255-020-0219-4

13. Weinberg F, Hamanaka R, Wheaton WW, et al. Mitochondrial metabolism and ROS generation are essential for Kras-mediated tumorigenicity. Proc Natl Acad Sci U S A. 2010;107(19):8788–8793. doi:10.1073/pnas.1003428107

14. Palaskas NJ, Garcia JD, Shirazi R, et al. Global alteration of T-lymphocyte metabolism by PD-L1 checkpoint involves a block of de novo nucleoside phosphate synthesis. Cell Discov. 2019;5(1). doi:10.1038/s41421-019-0130-x

15. Itzhaki O, Jacoby E, Nissani A, et al. Head-to-head comparison of in-house produced CD19 CAR-T cell in ALL and NHL patients. J Immunother Cancer. 2020;8(1):e000148. doi:10.1136/jitc-2019-000148

16. Rozenbaum M, Meir A, Aharony Y, et al. Gamma-Delta CAR-T Cells Show CAR-Directed and Independent Activity Against Leukemia. Front Immunol. 2020;11(July):1–8. doi:10.3389/fimmu.2020.01347

17. Jacoby E, Bielorai B, Hutt D, et al. Parameters of long-term response with CD28-based CD19 chimaeric antigen receptor-modified T cells in children and young adults with B-acute lymphoblastic leukaemia. Br J Haematol. 2022;197(4):475–481. doi:10.1111/bjh.18105

18. Sabatino M, Hu J, Sommariva M, et al. Generation of clinical-grade CD19-specific CAR-modified CD8 + memory stem cells for the treatment of human B-cell malignancies A platform for the generation of clinical-grade CD19-CAR modified T CD19-CAR modified T mediate superior antitumor responses com. Blood. 2016;128(4):519–529. doi:10.1182/blood-2015-11-683847

19. Klein Geltink RI, Edwards-Hicks J, Apostolova P, et al. Metabolic conditioning of CD8+ effector T cells for adoptive cell therapy. Nat Metab. 2020;2(8):703–716. doi:10.1038/s42255-020-0256-z

20. Sukumar M, Liu J, Ji Y, et al. Inhibiting glycolytic metabolism enhances CD8+ T cell memory and antitumor function. J Clin Invest. 2013;123(10):4479–4488. doi:10.1172/JCI69589

21. Wenes M, Jaccard A, Wyss T, et al. The mitochondrial pyruvate carrier regulates memory T cell differentiation and antitumor function. Cell Metab. 2022;34(5):731–746.e9. doi:10.1016/j.cmet.2022.03.013

22. Bai Z, Woodhouse S, Zhao Z, et al. Single-cell antigen-specific landscape of CAR T infusion product identifies determinants of CD19-positive relapse in patients with ALL. Sci Adv. 2022;8(23). doi:10.1126/sciadv.abj2820

23. Shiratori R, Furuichi K, Yamaguchi M, et al. Glycolytic suppression dramatically changes the intracellular metabolic profile of multiple cancer cell lines in a mitochondrial metabolism-dependent manner. Sci Rep. 2019;9(1):18699. doi:10.1038/s41598-019-55296-3

24. Aguer C, Gambarotta D, Mailloux RJ, et al. Galactose Enhances Oxidative Metabolism and Reveals Mitochondrial Dysfunction in Human Primary Muscle Cells. Luque RM, ed. PLoS One. 2011;6(12):e28536. doi:10.1371/journal.pone.0028536

25. van Bruggen JAC, Martens AWJ, Fraietta JA, et al. Chronic lymphocytic leukemia cells impair mitochondrial fitness in CD8+ T cells and impede CAR T cell efficacy. Blood. 2019;134(1):blood.2018885863. doi:10.1182/blood.2018885863

26. Weber EW, Parker KR, Sotillo E, et al. Transient rest restores functionality in exhausted CAR-T cells through epigenetic remodeling. Science. 2021;372(6537). doi:10.1126/science.aba1786

