## supplementary Fig. 1 for "Improved CAR-T cell activity associated with increased mitochondrial function primed by galactose"

**Supplementary File**

**Table of Content**

**Supplementary tables**

Supplementary table 1: Antibodies and dyes

Supplementary table 2: qPCR primers

**Supplementary figures**

Supplementary figure 1: Parameters of CAR-T cell products

Supplementary figure 2: CAR-T cell activation impacts mitochondrial parameters

Supplementary figure 3: Target cell growth in glucose and galactose

Supplementary figure 4: Growth of GluCARs and GalaCARs since PBMC activation

Supplementary figure 5: Activated GalaCARs have a greater capacity of increasing mitochondrial activity compared to GluCARs

Supplementary table 1: Antibodies and dyes

| **Antibody or dye** | **Brand** | **Clone** |
| --- | --- | --- |
| CAR reagent | Miltenyi Biotec |  |
| Anti-CD62L | Miltenyi Biotec | 145/15 |
| Anti-CD45RA | Miltenyi Biotec | REA562 |
| Anti-PD-1 | Miltenyi Biotec | PD1.3.1.3 |
| Anti-TIM-3 | R&D Systems | Rat IgG2A |
| GhostRed 780 Viability Dye | TONBO Biosciences |  |
| Anti-CD3 | Miltenyi Biotec | REA613 |
| Anti-CD4 | eBiosciences | RPA-T4 |
| Anti-CD8a | eBiosciences | SK1 |
| Anti-CD10 | Miltenyi Biotec | REA877 |
| Anti-CD20 | Miltenyi Biotec | REA780 |
| Streptavidin | BioLegend |  |
| Anti-4-1BB | BioLegend | 4B4-1 |
| Annexin dye | BioVision |  |
| CellTrace Violet Kit | Invitrogen |  |

Supplementary table 2: qPCR primers

| **Gene** | **Sequence (5’-3’)** |
| --- | --- |
| Human β2M (Fwd) | TGCTGTCTCCATGTTTGATGTATCT |
| Human β2M (Rev) | TCTCTGCTCCCCACCTCTAAGT |
| Human mitochondrial tRNA^leu^(Fwd) | CACCCAAGAACAGGGTTTGT |
| Human mitochondrial tRNA^leu^ (Rev) | TGGCCATGGGTATGTTGTTA |

β2M = β2-microglobulin

**Supplementary figures**

Supplementary figure 1:

Parameters of CAR-T cell products.

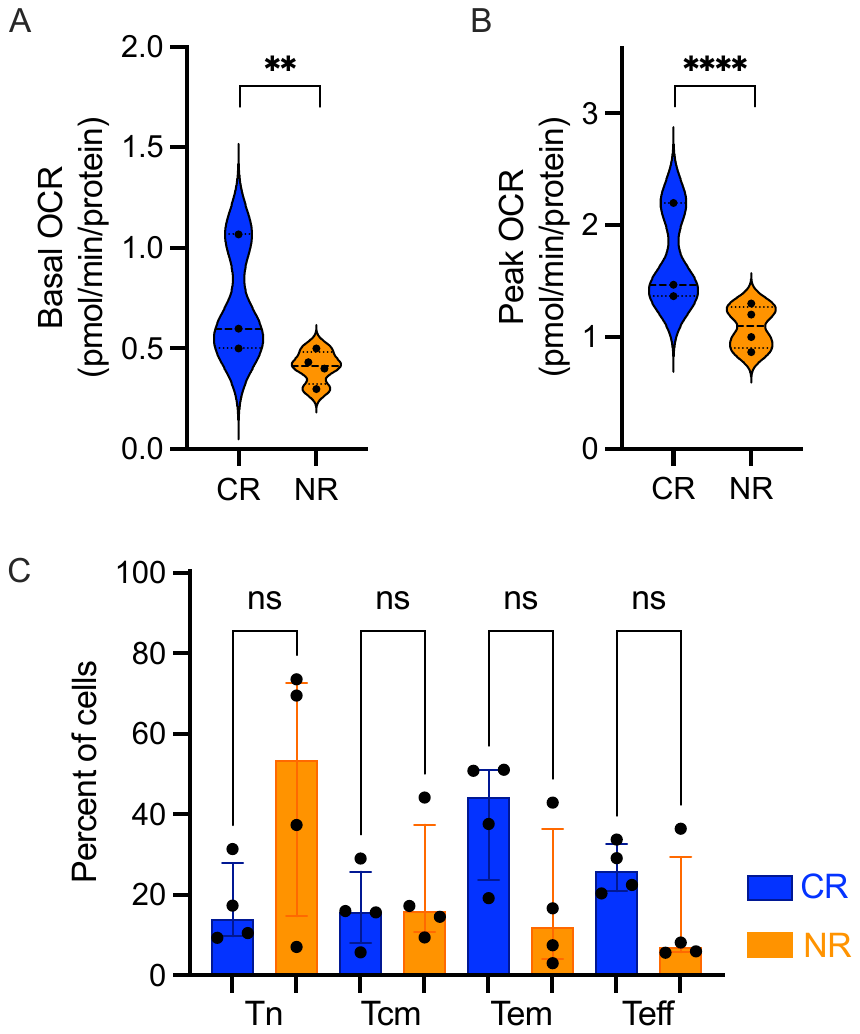

(A-B) Additional CAR-T products from patients with acute lymphoblastic leukemia were subjected to Oxygen Consumption Rate (OCR) measurements using the Seahorse XFe96 analyzer. (A) Basal mitochondrial OCR and (B) maximal respiration in products leading to response (CR, blue) or failing to achieve response (NR, non response, orange). (C) Thawed cultured products were subjected to flow cytometry analysis for memory subsets based on CD45RA and CD62L expression, and reported as naïve (Tn), central memory (Tcm), effector memory (Tem) and effector (Teff). Ns=non significant, **=p<0.01, **** p<0.0001.

Supplementary figure 2

CAR-T cell activation impacts mitochondrial parameters

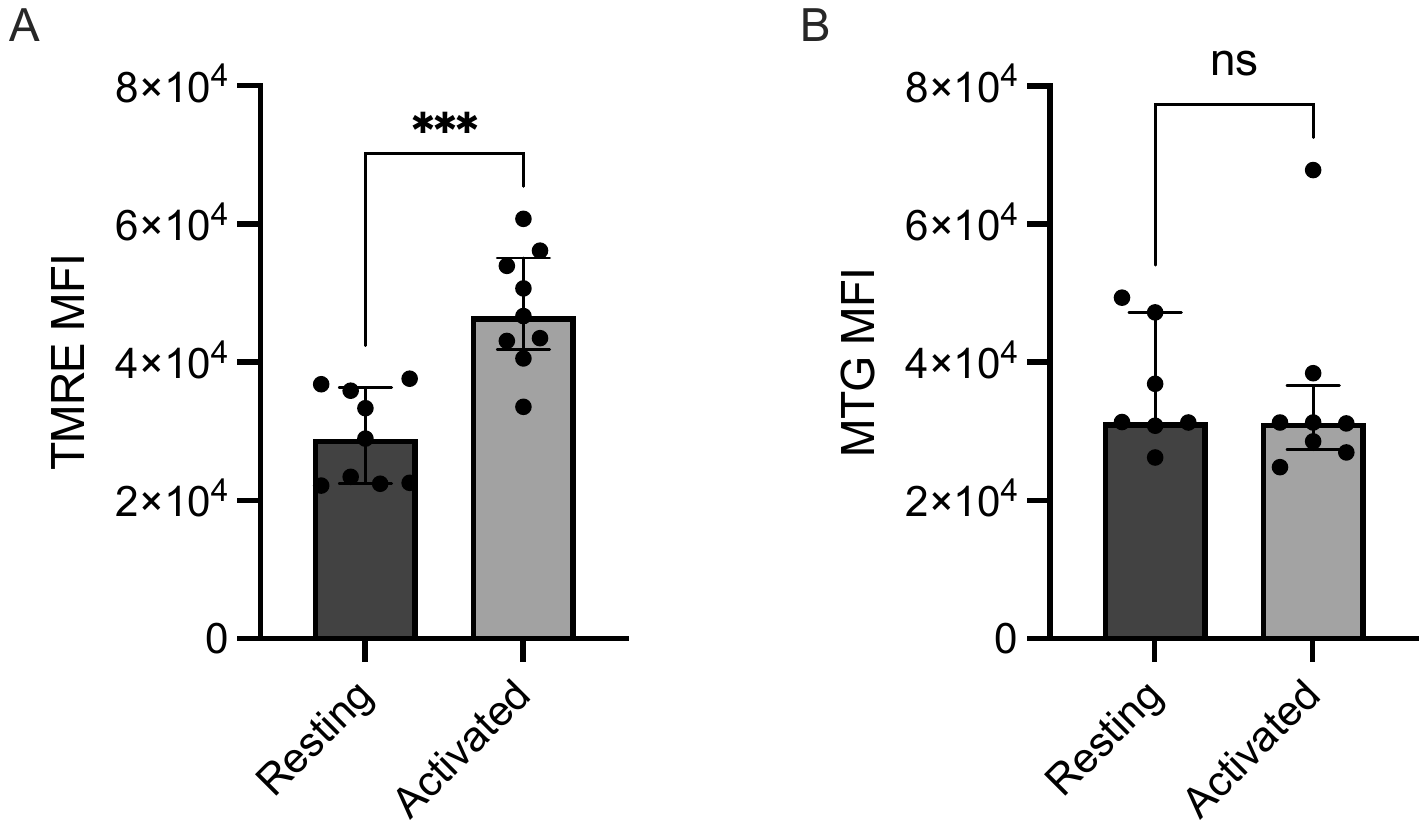

CAR-T cells either resting or activated using live or irradiated CD19+ cells were subjected to flow cytometry for tetramethylrhodamine methyl ester perchlorate (TMRE, A) to measure mitochondrial membrane potential, and MitoTracker Green (MTG, B) reflecting mitochondrial content. Mean fluorescent index of both dyes is shown. ***=p<0.001, ns=non significant.

Supplementary figure 3:

Target cell growth in glucose and galactose

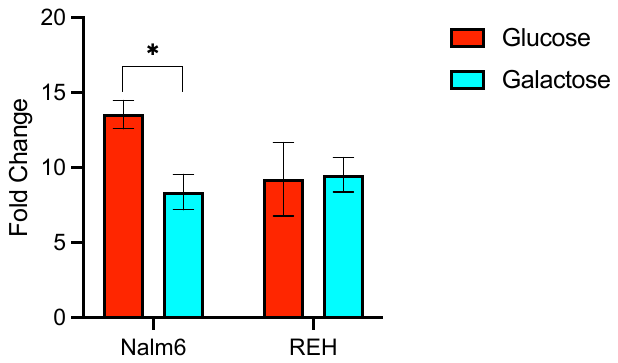

Two CD19+ cell lines were grown for 1 week in either glucose (red) or galactose (cyan). Nalm6 and REH are acute lymphoblastic leukemia cell lines. Growth fold change after 1 week is shown. *=p<0.05

Supplementary figure 4

Growth of GluCARs and GalaCARs since PBMC activation

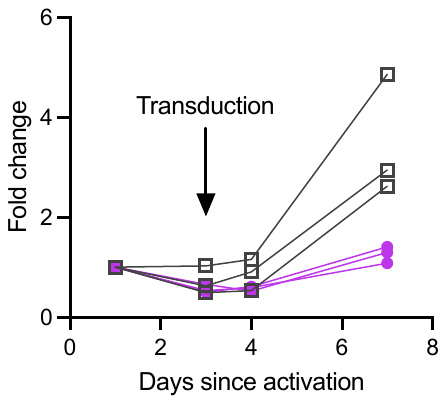

Growth of the T-cells in culture during production in either glucose (grey) or galactose (magenta) from initial peripheral blood mononuclear cell activation.

Supplementary figure 5

Activated GalaCARs have a greater capacity of increasing mitochondrial activity compared to GluCARs

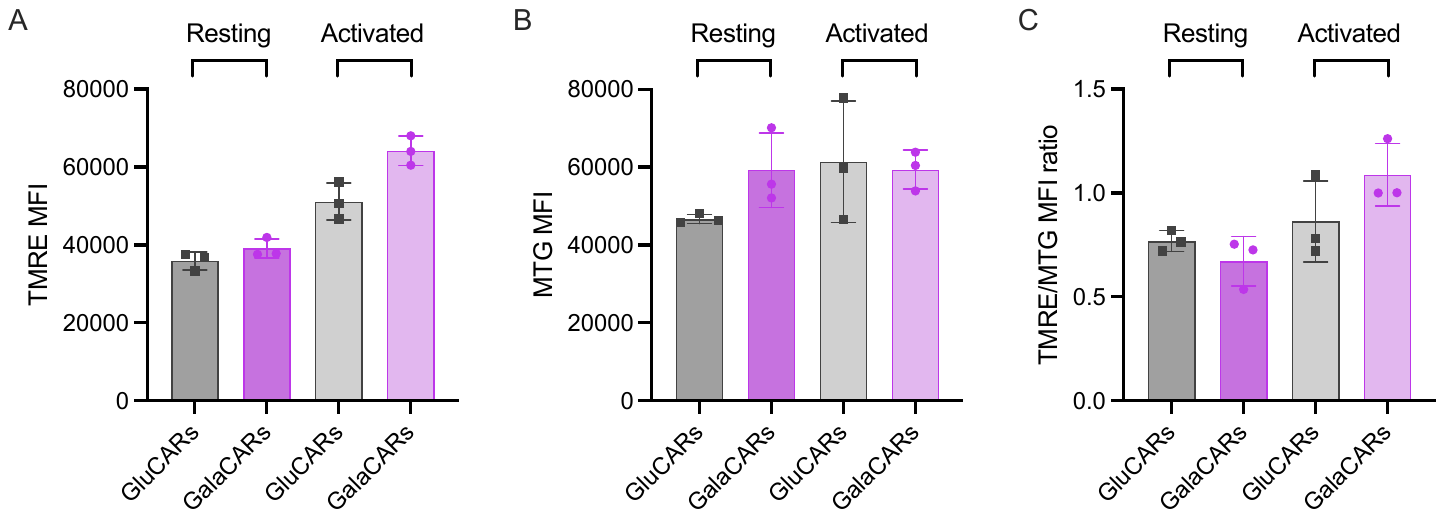

Resting and Nalm6-activated CAR-T cells grown in either standard glucose-based media (GluCARs) or galactose-based media (GalaCARs) were stained with TMRE or MTG followed by flow cytometry. Mean fluorescence index (MFI) of TMRE (A), MTG (B) and the ratio between the two (C) are shown in all conditions.
